# Chromosomal-level genome of fish mint *Houttuynia cordata* Thunb. (Saururaceae)

**DOI:** 10.1101/2024.04.25.590851

**Authors:** Sean T.S. Law, Wenyan Nong, Stacey S.K. Tsang, David T.W. Lau, Pang Chui Shaw, Jerome H.L. Hui

## Abstract

Herbaceous flowering plants in the family Saururaceae, or commonly known as the lizard’s tail family, are native to Southeast Asia and North America. Fish mint *Houttuynia cordata* is native to Southeast Asia and widely cultivated as culinary herb and medicinal plant in traditional medicine. Here, using a combination of PacBio HiFi long-read sequencing and Omni-C data, we present the chromosomal-level genome assembly of *H. cordata* (genome size 499.6 Mb). The genome has high sequence contiguity (scaffold N50 = 64.3 Mb) and completeness (BUSCO score of 94.6 %). 40,451 protein coding genes were also predicted using two transcriptomes generated in this study. The fish mint genome provides a valuable resource for further understanding the bioactive compounds and evolution of plants in the Saururaceae more widely.

## Background and Summary

The Saururaceae, or lizard-tail family, comprises of four genera (*Anemopsis, Gymnotheca, Houttuynia*, and *Saururus*) of herbaceous flowering plants native to Southeast Asia and North America. *Houttuynia cordata* Thunb., also commonly known as fish mint / chameleon plant / heart leaf / Chinese lizard plant / fish wort, is a perennial herb which belongs to the family Saururaceae (The Angiosperm Phylogeny Group [APG IV] 2016) (https://powo.science.kew.org/). This species is native to Southeast Asia with distributions in China, India, Indonesia, Japan, Korea, Myanmar, Nepal, Thailand and Vietnam (www.efloras.org) ^1,2^, and is characterized by its fishy smell, ovate-cordate leaves, creeping rhizomes, and a spike inflorescence with four white petaloid bracts resembling a single flower ^2^. It can be found in shady and moist wet location, and given its abilities to spread quickly and replace native plant species in woodland and wetland ecosystems ^34^, it has also been considered as invasive plants ^5^.

Other than its invasive property, its phytochemical properties have also been extensively studied. In many places in Asia, *H. cordata* has been widely cultivated as culinary herb and medicinal plant in traditional medicine ^6^. Its nutritional profile includes amino acids, crude proteins, lipids, vitamins (mainly vitamin-C), and carbohydrates; and depending on the geographical regions, either the edible rhizomes or leaves of *H. cordata* will be used in different cuisines ^7^. In addition *H. cordata* has also been widely used in traditional medicine such as Chinese, Indian, and Japanese in the last centuries, and various bioactive compounds such as alkaloids, flavonoids, fatty acids, polyphenols, organic acids, polysaccharides, sterols and volatile oils have also been identified ^7–16^.

Nevertheless, molecular resources of this ecological and cultural important *H. cordata* are limited to inter-simple sequence repeats ^17^, chloroplast genomes ^18–20^, and transcriptomes^21^, and the nuclear genomic resources remains limited, which hinders our understanding on their phytochemistry^22,23^ and evolutionary relationships ^24,25^. This study presents a high-quality chromosome-level genome of the fish mint *H. cordata*. The genome was assembled with a combination of PacBio HiFi long reads and Omni-C data. The resulting assembly is of 499.6 Mb in size with high contiguity and completeness, as indicated by a scaffold N50 value and BUSCO score of 64.3 Mb and 94.6%, respectively. Majority of the assembly (95.54%) was assigned into 11 pseudomolecules. Transcriptome sequencing from leaf and root tissues aided gene model prediction resulted into 40,451 protein coding genes with completeness BUSCO score of 87.6%. This high-quality *H. cordata* genome provides an essential resource for further investigations on their biology and ecology, such as the biosynthesis of phytochemicals as well as the evolutionary biology in Saururaceae.

## Methods

### Sample collection

Fresh samples of *H. cordata* were collected from the transplanted individuals in the Herbal Garden on the campus of The Chinese University of Hong Kong, which were originated from a local population in Ma On Shan, Hong Kong for more than 20 years.

### High molecular weight DNA extraction

High molecular weight (HMW) DNA was extracted from ∼0.5 g of leaf tissues using the Cetyltrimethylammonium bromide (CTAB) method (Doyle & Doyle, 1987) with slight modifications. Briefly, the tissues were ground into powder with liquid nitrogen and were transferred to 5 mL CTAB buffer containing 1% polyvinylpyrrolidone (PVP) and 0.2% 2-mercaptoethanol for 1 h digestion at 60°C. The lysate was centrifuged at 10,000 × *g* for 5 mins and the supernatant was transferred to a new 50 mL falcon tube. After treatment with the addition of 40 μL of RNase A for 5 mins, 1.6 mL 3M potassium acetate was added to the lysate and mixed by pipetting with a wide-bore tip. Upon 5 mins incubation at room temperature, the lysate were aliquoted into six 2 mL tubes, to which 800 μL of chloroform:isoamyl alcohol (24:1) was added separately and proceeded with the original protocol. The final HMW DNA was eluted with 60 µL elution buffer (PacBio Ref. No. 101-633-500) and was subjected to quality control with NanoDrop™ One Microvolume UV-Vis Spectrophotometer, Qubit® Fluorometer, and overnight pulse-field gel electrophoresis.

### Total RNA extraction and transcriptome sequencing

Total RNA was isolated from leaf and root tissues using mirVana miRNA Isolation Kit (Ambion) with CTAB pretreatment (Jordon-Thaden et al., 2015). The RNA samples were subjected to quality control using NanoDrop™ One Microvolume UV-Vis Spectrophotometer and gel electrophoresis. The samples were sent to Novogene for library construction and transcriptome sequencing (Table 1).

### PacBio library preparation and Hi-Fi sequencing

Approximately 5 μg of HMW DNA was sheared with a g-tube (Covaris Part No. 520079) for 6 rounds of centrifugation at 1,990 × *g* for 2 min, which was cleaned with SMRTbell® cleanup beads (PacBio Ref. No. 102158-300). Subsequently, a SMRTbell library was constructed with the SMRTbell® prep kit 3.0 (PacBio Ref. No. 102-141-700), following the manufacturer’s protocol. The library was eventually prepared with the Sequel^®^ II binding kit 3.2 (PacBio Ref. No. 102-194-100) and loaded with the diffusion loading mode with an on-plate concentration of 90 pM on the Pacific Biosciences SEQUEL IIe System using one SMRT cell. Finally, a 30-hour movie was run to output approximately 9.05 Gb of HiFi reads (Table 1).

### Omnic-C library preparation and sequencing

Plant nuclei was isolated from ∼2 g leaf tissues ground with liquid nitrogen using the protocol provided by PacBio (PacBio, 2022), which was modified from Workman et al. (2018). The nuclei pellet was resuspended with 4 mL 1X PBS buffer and proceeded with the Dovetail® Omni-C® Library Preparation Kit (Dovetail Cat. No. 21005), following the manufacturer’s instructions. The resulting library was subject to quality control using Qubit^®^ Fluorometer and TapeStation D5000 ScreenTape for concentration and fragment size measurement, respectively. The final Omni-C library was then sent to Novogene and sequenced on the Illumina HiSeq-PE150 platform, from which approximately 33.57 Gb data were generated (Table 1).

### Genome assembly and genome characteristics estimation

*De novo* genome assembly was performed using Hifiasm (version 0.16.1-r375) with default parameters (Cheng et al., 2021)^26^. Haplotypic duplications were identified and removed using purge_dups based on the depth of HiFi reads (Guan et al., 2020)^27^. Proximity ligation data from the Omni-C library were used to scaffold genome assembly using YaHS (version 1.2a.2) with default parameters (Zhou et al., 2023)^28^. Benchmarking Universal Single-Copy Orthologs (BUSCO, v5.5.0) (Manni et al., 2021)^29^ was run with the Viridiplantae dataset (Viridiplantae Odb10) to assess the completeness of the assembly. Regarding the genome characteristics of the assembly, the k-mer count and histogram were generated at k = 21 from Omni-C reads using Jellyfish (v2.3.0) (Marçais & Kingsford, 2011)^30^ with the parameters “count -C -m 21 -s 1000000000 -t 10”, and the reads.histo was uploaded to GenomeScope to estimate genome heterozygosity, repeat content and size using default parameters (v2.0) (http://qb.cshl.edu/genomescope/genomescope2.0/) (Ranallo-Benavidez et al., 2020)^31^. The final assembly resulted in a total genome size of 499.6 Mb, with a BUSCO score and scaffold N50 of 94.6% and 64.3 Mb, respectively (Table 2; Figure 1B). Majority (95.54%) of the scaffolds were anchored into 11 pseudomolecules (Table 3; Figure 1C), which correspond to the range of basic chromosome number of *H. cordata* (Luo et al., 2022)^32^. Genome characteristic estimation resulted an estimated genome size of 549.3 Mb with a low rate of heterozygosity at 0.96% (Figure 1C).

**Figure 1.**
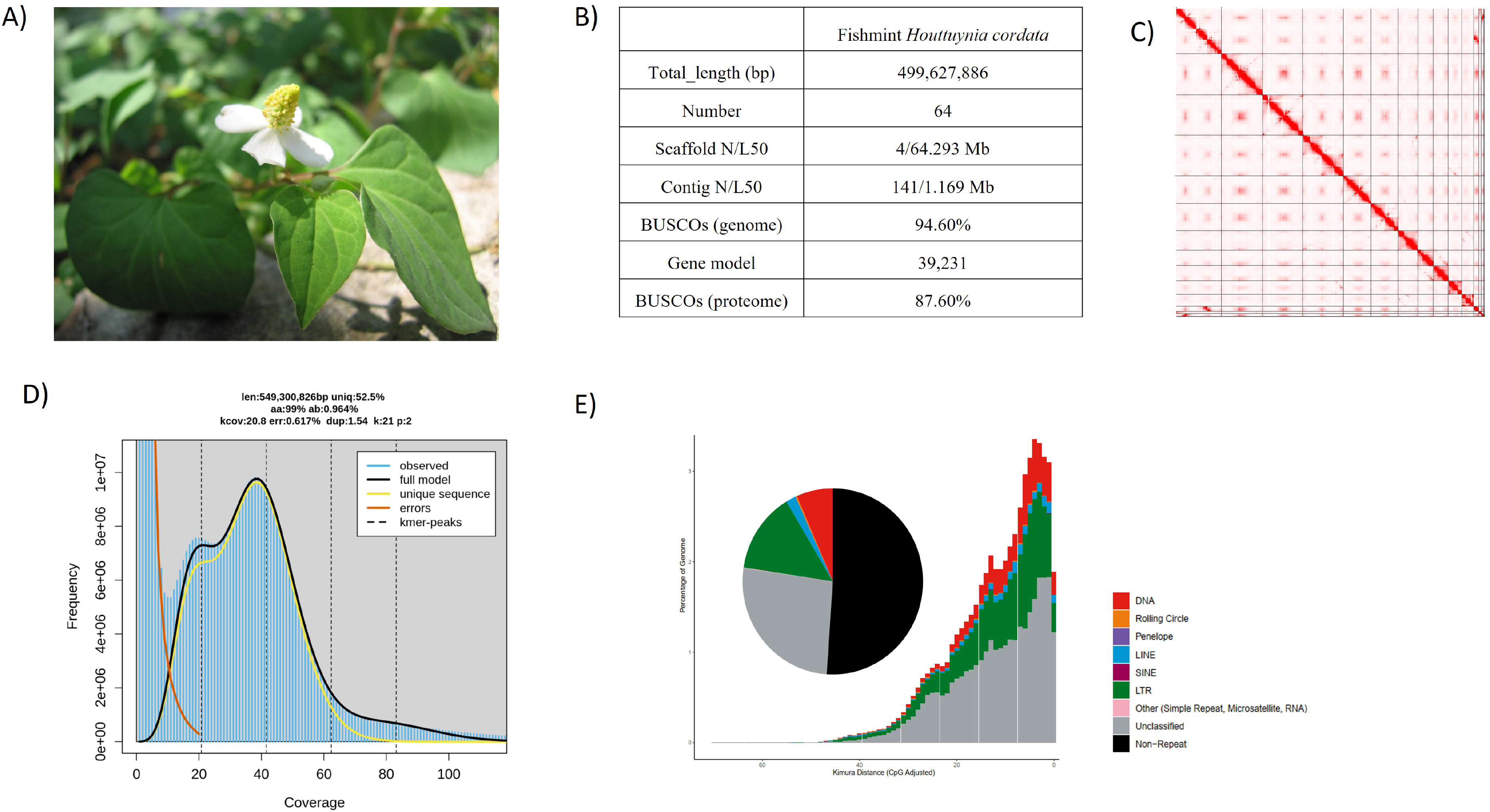
A) Picture of *H. cordata*; B) Statistics of the genome assembly generated in this study; C) Hi-C contact map of the assembly; D) Estimated genome size (*K*-mer = 21); E) Repetitive elements distribution.

### Gene model prediction

Gene models were predicted by funannotate (version 1.8.15) (Palmer & Stajich, 2020)^33^ using the parameters “--protein_evidence uniprot_sprot.fast --cpus 50 -- genemark_mode ET --optimize_augustus --busco_db embryophyta --organism other -- max_intronlen 350000 --name Hcor”. Briefly, the genome was soft-masked by redmask (v0.0.2) (https://github.com/nextgenusfs/redmask) (Girgis et al., 2015)^34^. Raw RNA sequencing data were first processed using Trimmomatic (version 0.39) with the parameters “TruSeq3-PE.fa:2:30:10 SLIDINGWINDOW:4:5 LEADING:5 TRAILING:5 MINLEN:25” (Bolger, Lohse & Usadel, 2014)^35^ to remove the low quality reads, and using kraken2 (v2.0.8 with kraken2 database k2_standard_20210517) (Wood, Lu & Langmead, 2019)^36^ to remove the contaminated reads. The processed reads were assembled by Trinity (v2.8.5) with parameters “--stranded RF --max_intronlen 350000” (Grabherr MG et al. 2011)^37^, the Trinity transcript was used to map to the repeat soft-masked genome by minimap2 (v2.2.1) with default parameters (Li, 2018)^38^. The Trinity transcript alignments were converted to GFF3 format and used as input to run the PASA alignment (v2. 5.3) with default parameters (Haas et al., 2008)^39^ in the Launch_PASA_pipeline.pl process to obtain the PASA models trained by TransDecoder (v5.7.1) with default parameters (https://github.com/TransDecoder/TransDecoder), and then using Kallisto TPM data (v 0.46.1) to select the PASA gene models. The PASA gene models were used to train Augustus in the funannotate-predict step. The gene models from several prediction sources, including GeneMark (v3.68_lic) (Lomsadze et al., 2014)^40^, high-quality Augustus predictions (HiQ), PASA (v2.5.3) (Haas et al., 2008)^39^, Augustus (v3. 5.0) (Stanke et al., 2006)^41^, GlimmerHM (v 3.0.4) (Majoros et al. 2004)^42^ and snap (v2006-07-28) (Korf 2004)^43^ were passed to Evidence Modeler (v1.1.1) with EVM Weights {‘Augustus’: 1, ‘HiQ’: 2, ‘GeneMark’: 1, ‘GlimmerHMM’: 1, ‘pasa’: 6, ‘snap’: 1, ‘proteins’: 1, ‘transcripts’: 1} to generate the gene model annotation files and the UTRs were captured in the funannotate-update step using PASA. BUSCO (v5.5.0) (Manni et al., 2021)^29^ was run with the Viridiplantae dataset (Viridiplantae Odb10) to assess the completeness of gene model dataset. A total of 39,231 gene models with a total of 41,195 transcripts were generated in the final genome annotation files, including 744 tRNA and 40,451 protein-coding genes, with a BUSCO score of 87.6% (Table 2; Figure 1B).

### Repeat annotation

Transposable elements (TEs) were annotated as previously described (Baril et al, 2022)^44^ using the automated Earl Grey TE annotation pipeline (version 1.2, https://github.com/TobyBaril/EarlGrey) with “-r eukarya” to search the initial mask of known elements and other default parameters. Briefly, this pipeline first identified known TEs from Dfam with RBRM (release 3.2) and RepBase (v20181026). De novo TEs were then identified, and consensus boundaries extended using an automated BLAST, Extract, Extend process with 5 iterations and 1000 flanking bases added in each round. Redundant sequences were removed from the consensus library before the genome assembly was annotated with the combined known and de novo TE libraries. Annotations were processed to remove overlap and defragment annotations prior to final TE quantification. Eventually, 244.4 Mb of TEs were annotated, comprising about half (48.9%) of the fish mint genome (Table 4; Figure 1E). The major classified TEs were primarily LTR retrotransposons (14.0%), DNA transposons (6.52%) and LINE transposons (1.81%).

## Data Records

The raw reads generated in this study, including Transcriptome (SAMN40408130 and SAMN40408131), Omni-C (SRR28296043) and PacBio HiFi (SRR28296042) data, have been deposited in the NCBI database under the BioProject accession number PRJNA1086680 (https://www.ncbi.nlm.nih.gov/bioproject/PRJNA1086680). The genome, genome and repeat annotation files have been deposited and are publicly available in figshare (https://figshare.com/s/bade9d25b6338655e7ec).

## Technical Validation

The pseudochromosomes of the final assembly were validated by inspecting the Omni-C contact maps using Juicer tools (version 1.22.01) (Durand et al., 2016). Briefly, Omni-C reads were mapped and aligned by BWA with parameters “mem -5SP -T0”, the parsing module of the pairtools pipeline was used to find ligation junctions with parameters “--min-mapq 40 --walks-policy 5unique --max-inter-align-gap 30 --nproc-in 8 --nproc-out 8”. The parsed pairs were then sorted using pairtools sort with default parameters, PCR duplicate pairs were removed using pairtools dedup with parameters “--nproc-in 8 --nproc-out 8 -- mark-dups”, the pairs file was split using pairtools split with default parameters and used to generate the contact matrix using juicertools and Juicebox (v1.11.08).

## Code availability

biopython: 1.81; goatools: 1.3.1; matplotlib: 3.4.3; natsort: 8.4.0; numpy: 1.24.4; pandas: 1.5.3; psutil: 5.9.5; requests: 2.31.0; scikit-learn: 1.3.0; scipy: 1.10.1; seaborn: 0.12.2; PASA: 2.5.3; CodingQuarry: 2.0; Trinity: 2.8.5; augustus: 3.5.0; bamtools: bamtools 2.5.1; bedtools: bedtools v2.31.0; blat: BLAT v37x1; diamond: 2.1.8; emapper.py: 2.1.12; ete3: 3.1.3; exonerate: exonerate 2.4.0; fasta: 36.3.8g; glimmerhmm: 3.0.4; gmap: 2021-12-17; gmes_petap.pl: 4.71_lic; hisat2: 2.2.1; hmmscan: HMMER 3.3.2 (Nov 2020); hmmsearch: HMMER 3.3.2 (Nov 2020); java: 17.0.3-internal; kallisto: 0.46.1; mafft: v7.520 (2023/Mar/22); makeblastdb: makeblastdb 2.14.1+; minimap2: 2.26-r1175; pigz: 2.6; proteinortho: 6.3.0; salmon: salmon 0.14.1; samtools: samtools 1.16.1; signalp: 5.0b; snap: 2006-07-28; stringtie: 2.2.1; tRNAscan-SE: 2.0.12 (Nov 2022); tantan: tantan 40; tbl2asn: 25.8; tblastn: tblastn 2.14.1+; trimal: trimAl v1.4.rev15 build[2013-12-17]; trimmomatic: 0.39

## Authors Contributions

JHLH and PCS conceived and supervised the study. DTWL, PCS, JHLH acquired funding. SSKT collected samples and culture them in the laboratory. STSL and SSKT extracted DNA and RNA, and constructed PacBio and Omni-C libraries for sequencing. WN assembled and genome and carried out gene model prediction. STSL and WN carried out the validation.

## Acknowledgements

This work was supported by Li Dak Sum Yip Yio Chin R & D Centre for Chinese Medicine, Innovation Technology Fund of Innovation Technology Commission: Funding Support to State Key Laboratory of Agrobiotechnology (8300040), and Direct Grant of The Chinese University of Hong Kong.

## Figure and Table legend

**Table 1**. Genome and transcriptome sequencing data.

**Table 2**. Genome assembly information.

**Table 3**. BUSCO scores.

**Table 4**. Repeat content summary.

**Supplementary Information 1**. Genome assembly quality control (QC) and contaminants detection.

